# Introducing a New Bond-Forming Activity in an Archaeal DNA Polymerase by Structure-Guided Enzyme Redesign

**DOI:** 10.1101/2021.12.28.474375

**Authors:** Tushar Aggarwal, William A. Hansen, Jonathan Hong, Abir Ganguly, Darrin M. York, Sagar D. Khare, Enver Cagri Izgu

## Abstract

DNA polymerases have evolved to feature a highly conserved activity across the tree of life: formation of, without exception, phosphodiester linkages that create the repeating sugarphosphate backbone of DNA. Can this linkage selectivity observed in nature be overcome by design to produce non-natural nucleic acids? Here, we report that structure-guided redesign of an archaeal DNA polymerase (9°N) enables a new polymerase activity that is undetectable in the wild type enzyme: catalyzing the formation of N3’→P5’ phosphoramidate linkages in the presence of 3’-amino-2’,3’-dideoxynucleoside 5’-triphosphate (3’-NH_2_-ddNTP) building blocks. Replacing a highly conserved metal-binding aspartate in the 9°N active site (Asp-404) with asparagine was key to the emergence of this unnatural enzyme activity. Molecular dynamics simulations provided insights into how a single substitution could enhance the productive positioning of the 3’-amino nucleophile in the active site. Further remodeling of the protein-nucleic acid interface with substitutions in the finger subdomain led to a quadruple-mutant variant (9°N-NRQS) that incorporated 3’-NH_2_-ddNTPs into a 3’-amino-primer on various DNA templates. This work presents the first example of an active-site substitution of a metal-binding residue that leads to a novel activity in a DNA polymerase, and sheds light on the molecular basis of substrate fidelity and latent promiscuity in enzymes.

## INTRODUCTION

Transfer of genetic information is orchestrated by protein enzymes in all extant organisms on Earth. DNA replication, the process of producing copies of DNA, is essential for information transfer and catalyzed by diverse DNA polymerases. Despite large differences in composition, catalytic activity, and replication fidelity, all known DNA polymerases have evolved to feature a stringent internucleotidyl linkage selectivity.^1^ They catalyze the formation of O3’→P5’ phosphodiester bond between the 3’-OH of a DNA primer and the 5’-a-phosphate of a 2’-deoxynucleoside triphosphate (dNTP) monomer, leading to the repeating sugar-phosphate backbone of DNA. This transformation requires the presence of divalent metal cations, typically Mg^2+^, which activate the 3’-OH nucleophile and thereby reduce the activation energy to reach the phosphorylation transition state.^2,3^

Despite exquisite preference in utilizing hydroxyl nucleophiles of dNTPs, certain DNA polymerases can display low level promiscuous activities. For example, they can accept non-canonical nucleotides with modifications in the nucleobase^4,5^ or sugar^6–10^ framework and synthesize non-natural nucleic acids.^11^ However, enzymatic formation of sugar-phosphate linkages between atoms other than oxygen and phosphorous has rarely been observed, suggesting that the molecular machinery essential for genetic information transfer has evolved to favor, with high stringency, O-P bond formation.

One distinct nucleic acid that contains a non-O–P internucleotidyl linkage is 3’-nitrogen-5’-phosphorus-linked DNA (NP-DNA), where each 3’-O atom is replaced by a 3’-NH, thus possessing N3→P5’ phosphoramidate linkages (**Figure 1**). NP-DNA mimics RNA in that its duplex structure adopts an RNA-like A-form helical geometry in solution and *in crystallo*^12^ Notably, reverse transcriptases can recognize templates made of NP-DNA and synthesize a complementary DNA strand.^13^ Additionally, NP-DNA can form duplexes with both ssDNA and ssRNA that exhibit higher thermal stability than isosequential duplexes made of DNA and RNA.^12^ Furthermore, NP-DNA has been proposed as a potential inhibitor of RNAs and RNA-binding proteins.^14,15^ NP-DNA can inhibit gene expression both *in vitro* and *in vivo* by acting as a steric block in translation without the need to promote endogenous RNase H activity.^16,17^ Thus, NP-DNA has chemical and structural features that make it a unique polymer with significant potential in synthetic biology and biotechnology.

**Figure 1.**
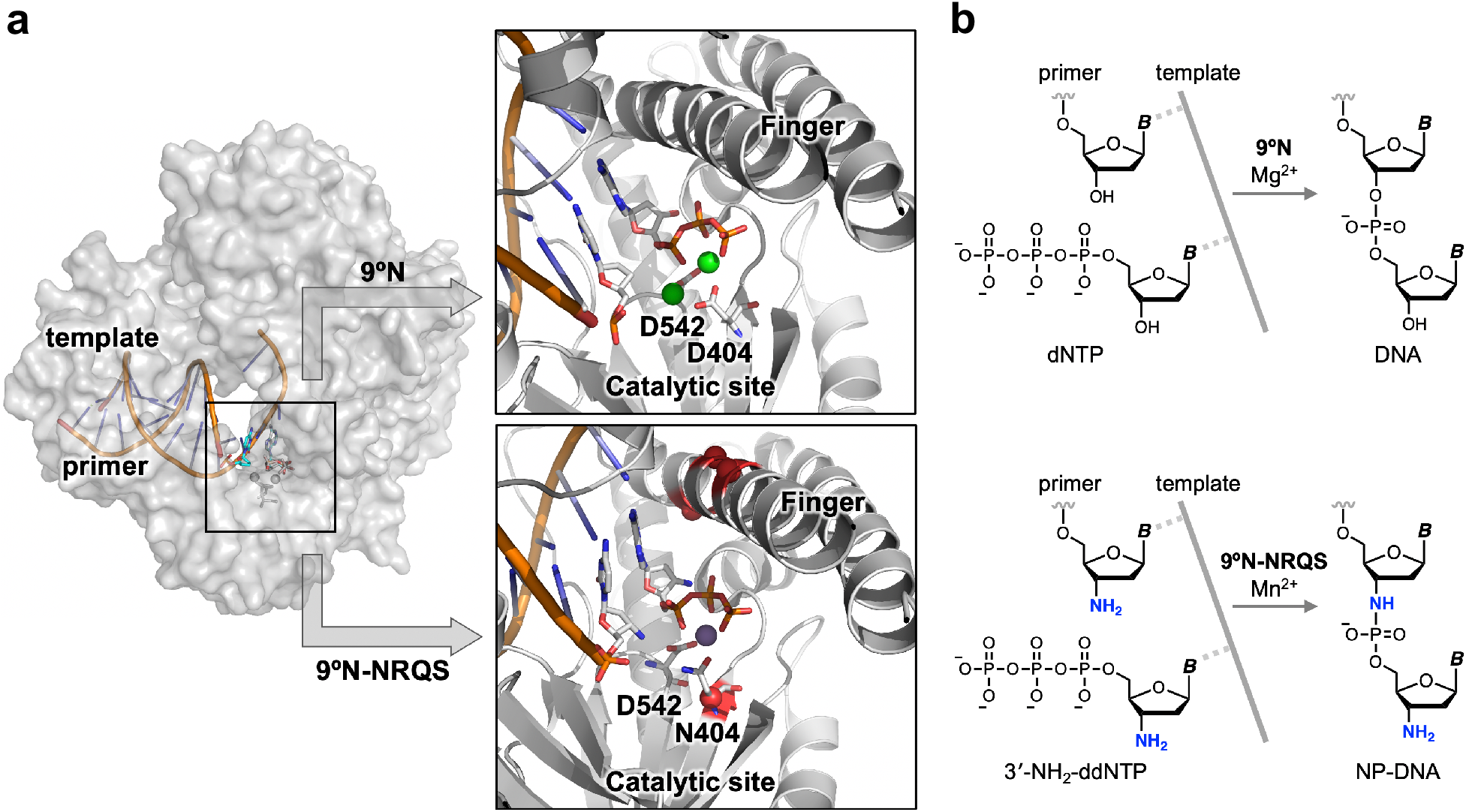
Structure-guided redesign of 9°N DNA polymerase to catalyze the formation of internucleotidyl N-P bonds. **(a)** Model of 9°N bound to a primer/template complex. Insets show the positioning of the primer and the incoming nucleoside triphosphate within the WT 9°N (top, PDB accession code: 5OMV) and its quadruple-mutant variant 9°N-NRQS (bottom). Sites of substitutions in 9°N-NRQS are highlighted by red spheres. Active site metal ions are shown: Mg^2+^ in green (top) and Mn^2+^ in purple (bottom). **(b)** Template-directed primer-extension reactions catalyzed by WT 9°N (top) and 9°N-NRQS (bottom).

Enzymatic synthesis of NP-DNA is fundamentally challenging, as phosphoramidate bond formation is not a primary function of known wild-type (WT) polymerases. However, a WT DNA polymerase from *Bacillus stearothermophilus* has recently been reported to display low level promiscuous N3→P5’ bond formation activity.^18^ Structural characterizations indicated single-metal ion coordination to nucleoside triphosphate (in the B-site), but no coordination to the primer 3’-amino group (in the A-site). Indeed, in the absence of divalent metal ions, nonenzymatic nucleic acid primer phosphorylation is substantially faster with a 3’-amino nucleophile compared to 3’-hyrdoxyl nucleophile.^19^ In contrast, at least two divalent metal ions are needed in polymerase-catalyzed canonical primer phosphorylation.^2,3^ We therefore hypothesized that a WT DNA polymerase that has undetectable activity in catalyzing N3’→P5’ bond formation could be redesigned with ablated A-site metal occupancy to exhibit this novel function (**Figure 1**). Although certain residues in the active site of WT DNA polymerases can be modified without major impact on their activity, metal-binding Asp residues are considered immutable without complete loss of function.^20^ Consequently, we reasoned that substituting these metal-binding residues could result in an extreme trade-off between the WT and novel internucleotidyl linkage selectivities, which is rare in enzyme evolution but not uncommon in designed biocatalysts.^21,22^

In this work, using structure-guided computational design, we engineered variants of 9°N (**Figure 1**), an archaeal B-family replicative DNA polymerase, to synthesize NP-DNA oligomers from 3’-amino-2’,3’-dideoxynucleoside 5’-triphosphates (3’-NH2-ddNTPs). Replacing metal-binding Asp-404 in the 9°N active site with asparagine (Asn) dramatically decreased its DNA polymerase activity. Strikingly, however, this substitution provided the desired novel function: catalyzing N3→P5’ phosphoramidate bond formation. Computational modeling revealed that productive nucleophile positioning in the active site was key for the emergence of the novel activity. The polymerase processivity was enhanced via three amino acid substitutions in the distal subdomains of the polymerase. The resulting quadruple-mutant variant displayed DNA-dependent NP-DNA polymerase activity with modest rates and fidelity. This work, to our knowledge, presents the first example of substitution of metal-binding amino acids in a DNA polymerase active site leading to the catalysis of phosphoramidate bond formation, endowing a novel function to archaeal B-family DNA polymerases.

## RESULTS

### Polymerase Active-Site Remodeling for Catalyzing Phosphoramidate Bond Formation

9°N displays intrinsic thermostability,^23^ can tolerate amino acid mutations,^24^ and can accommodate non-canonical nucleoside triphosphates as monomers in synthesis of RNA-like nucleic acids.^6,7^ Given these versatile characteristics, we selected 9°N as the starting point in our investigation of the requirements for enzymatic internucleotidyl N3→P5’ phosphoramidate bond formation.

Divalent metal cations can affect the rate and fidelity of nucleic acid synthesis catalyzed by DNA polymerases.^25^ However, there is no established evidence to suggest that 9°N would operate under the same conditions to form phosphoramidate bonds for a 3’-amino primer. A DNA polymerase from *Bacillus stearothermophilus* has recently been reported to favor a single-metal ion coordination that primarily affects B-site metal ion, but not the A-site metal ion during the catalysis of phosphoramidate bond formation.^18^ To investigate the metaldependence of 9°N in extending a 3’-amino primer on a DNA template (**Table S1**), we screened various divalent metal ions, including Mg^2+^, Ca^2+^, Mn^2+^, Zn^2+^, Ni^2+^, Co^2+^, Fe^2+^, and Cu^2+^ (**Figure S1**). We set the pH of these and the other primer extension reaction mixtures in this work as 8.8. At this pH, a substantial population of the 3’-amino nucleophile is expected to be neutral, as the *pK_a_* of the 3’-ammonium group of 2’,3’-dideoxynucleosides is ~7.5–7.7.^19,26^ We used 3’-NH2-ddTTP (1 mM) as the monomer and a primer/template complex composed of a 5’-FAM labeled 15-nt DNA primer ending with a 3’-NH2-G (1 μM) and a 22-nt complementary DNA template containing a 5’-C2A5 overhang (1 μM) (**Table S1**). Denaturing gel analysis indicated that 9°N (1 μM) failed to extend the 3’-amino primer at 55 °C with any of these divalent metals (**Figure S1**), raising the possibility that they may hinder the nucleophilic character of the 3’-amino group within the 9°N active site. Therefore, we hypothesized that decreasing metal ion occupancy around the primer 3’-amino group could facilitate a more productive 9°N-primer interaction vis-à-vis the formation of phosphoramidate bonds.

9°N possesses two conserved Asp residues in the active site, Asp-404 and Asp-542, each of which coordinates a divalent metal cation.^27^ Currently, there is no report showing which of these two 9°N Asp residues coordinates to the metal ion that is expected to occupy the space near a primer 3’-amino group. Therefore, we synthesized a total of five 9°N Asp variants (**Table S2** and **Figure S2A**), each constituting a variation in side chain polarizability^28^: Asn, Ala, Ser, His, and Thr. For one of the five variants, Asp-404 was substituted with Asn (9°N-N), and for the other four, Asp-542 was substituted with either Ala (9°N-A’), Ser (9°N-S), Thr (9°N-T), or His (9°N-H). Rosetta-based models of these variants showed steric compatibility with the wild type enzyme conformation, suggesting that all of the variants should be sufficiently stable. Indeed, we were able to produce these proteins by recombinant expression (**Figure S2**). However, under the reaction conditions described above, N3→P5’ phosphoramidate bond formation (+1-nt product) was observed primarily for the polymerase-metal ion combination of 9°N-N and Mn^2+^ (**Figure S1**). These results indicate that low-level phosphoramidate bond formation is a latent promiscuous activity of the polymerase that can be accessed by changing merely a single non-hydrogen atom (oxygen to nitrogen) in a sidechain group.

### Molecular Dynamics Simulations to Investigate the Consequences of Asp-to-Asn Substitution and Role of Mn^2+^

To gain a better understanding of the influence of D404N substitution, A-site metal ion, and the identity of primer 3’ nucleophile on the structure and dynamics of the active site of 9°N, we performed exhaustive molecular dynamics (MD) simulations of 9°N with various active site configurations. In total, we carried out ~0.5 μs-long, independent MD simulations for 8 different active site configurations of 9°N, specifically the WT and D404N variant, in presence of either a 3’-OH or 3’-NH_2_ primer, in presence and absence of the A-site metal ion. The results from these MD simulations are summarized in **Figure 2** as 2D scatter plots of the in-line attack angle (τ, the angle between O3’, scissile phosphorus atom, and O5’) and the distance between the 3’-nucleophile and scissile phosphorus atom (r), created from the corresponding MD configurations. The configurations are classified as *active (A)* or *non-active (NA),* where *active* configurations are defined as those in which the active site possess in-line fitness (t > 150°) and in which the 3’-nucleophile is favorably positioned for catalysis (distance between 3’ nucleophile and P ≤ 3.5 Å).

**Figure 2.**
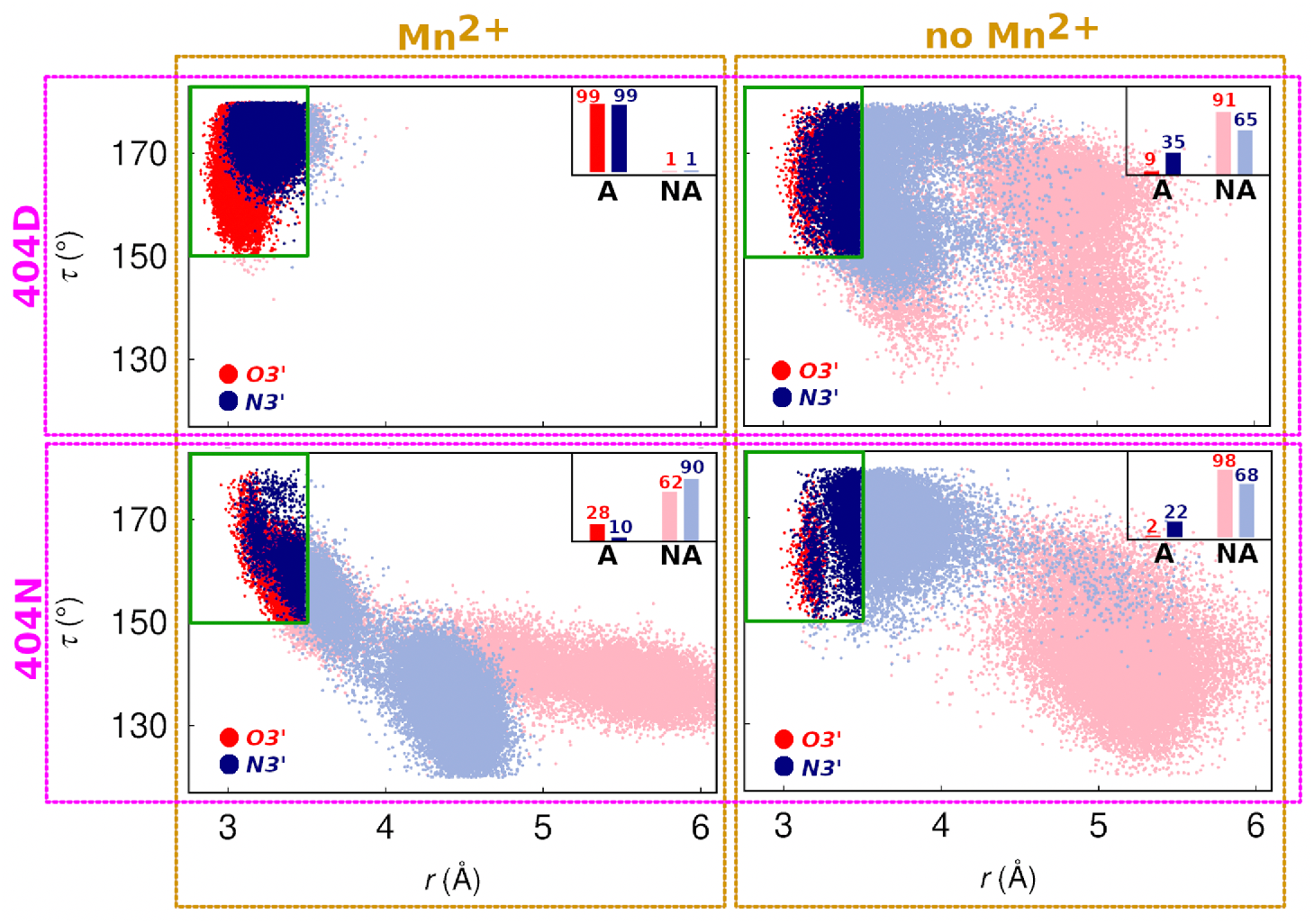
Analysis of 3’-nucleophile positioning using Molecular Dynamics (MD) simulations. 2D scatter plots of the in-line attack angle (τ, the angle between the 3’ nucleophile, scissile phosphorus atom, and O5’) and the distance between the 3’ nucleophile and scissile phosphorus atom (r), obtained from MD simulations of 9°N in various active site configurations. In total, eight different active site scenarios are explored: the four panels correspond to simulations in which residue 404 is ASP and A-site metal ion is present (top left), residue 404 is ASP and A-site metal ion is absent (top right), residue 404 is ASN and A-site metal ion is present (bottom left), and residue 404 is ASN and A-site metal ion is absent (bottom right). In each panel, the red and blue colors correspond to the primer being 3’-OH (red) or 3’-NH_2_ (blue). The data points are classified as *active (A)* or *non-active (NA),* where *active* configurations are defined as those in which the active site possesses in-line fitness (t > 150°) and in which the 3’ nucleophile is favorably positioned for catalysis (distance between 3’ nucleophile and P ≤ 3.5 Å).

#### WT Polymerase Simulations

In the WT polymerase ensemble, 3’-OH is favorably positioned for catalysis in >90% of Asp-404 configurations when Mn^2+^ is present (**Figure 2**, top left, red), compared to a 10-fold decrease (9%) in active configurations when Mn^2+^ is absent (**Figure 2**, top right, red), suggesting that the presence of divalent metals is crucial to the reactivity of primer 3’-OH in 9°N. Similarly, primer 3’-NH2 is favorably positioned for catalysis in >90% of Asp-404 configurations when Mn^2+^ is present, with a corresponding 3-fold decrease (35%) in active configurations when Mn^2+^ is absent (**Figure 2**, top left, blue), indicating that there is still a significant fraction of active Asp-404 configurations without Mn^2+^. These simulations provide further support to our hypothesis that coordination of the 3’-amino primer to a divalent metal is unproductive for primer extension.

#### D404N Variant Polymerase Simulations

3’-OH was favorably positioned in 28% of Asn-404 configurations when Mn^2+^ was present (**Figure 2**, bottom left, red), compared to a 14-fold decrease (2%) in active configurations when the Mn^2+^ was absent (**Figure 2**, bottom right, red). In contrast, 3’-NH2 was favorably positioned for catalysis in 10% of Asn-404 configurations when Mn^2+^ was present (**Figure 2**, bottom left, blue), compared to a ~2-fold increase (22%) when Mn^2+^ was absent (**Figure 2**, bottom right, blue). Taken together, these simulations support a scenario in which i) positioning of both 3’-OH and 3’-NH2 are influenced by both the identity of the 404 residue and the presence of Mn^2+^, ii) phosphoramidate bond formation may occur in the absence of Mn^2+^ near the primer 3’-amino group, and iii) the primer 3’-amino group may act as a more potent nucleophile in the absence of Mn^2+^.

### Investigating the Role of Active Site and Finger Domains in Affecting the Rate of Phosphoramidate Bond Formation via Structure-guided Mutagenesis

Using Rosetta-based modeling,^29^ we explored additional substitutions beyond D404N throughout 9°N (**Figure 3a**). With these 9°N variants (**Figure S2** and **Table S2**), we conducted a comparative polymerase activity analysis for the extension of the same 3’-amino primer on a 5’-A_2_C_5_ DNA template with 3’-NH_2_-ddGTP at 55 °C (**Figure S3**). We measured the observed rate constant, *k*_obs_, for the primer extension by following the depletion of the primer over 24 h (**Table S4**).

**Figure 3.**
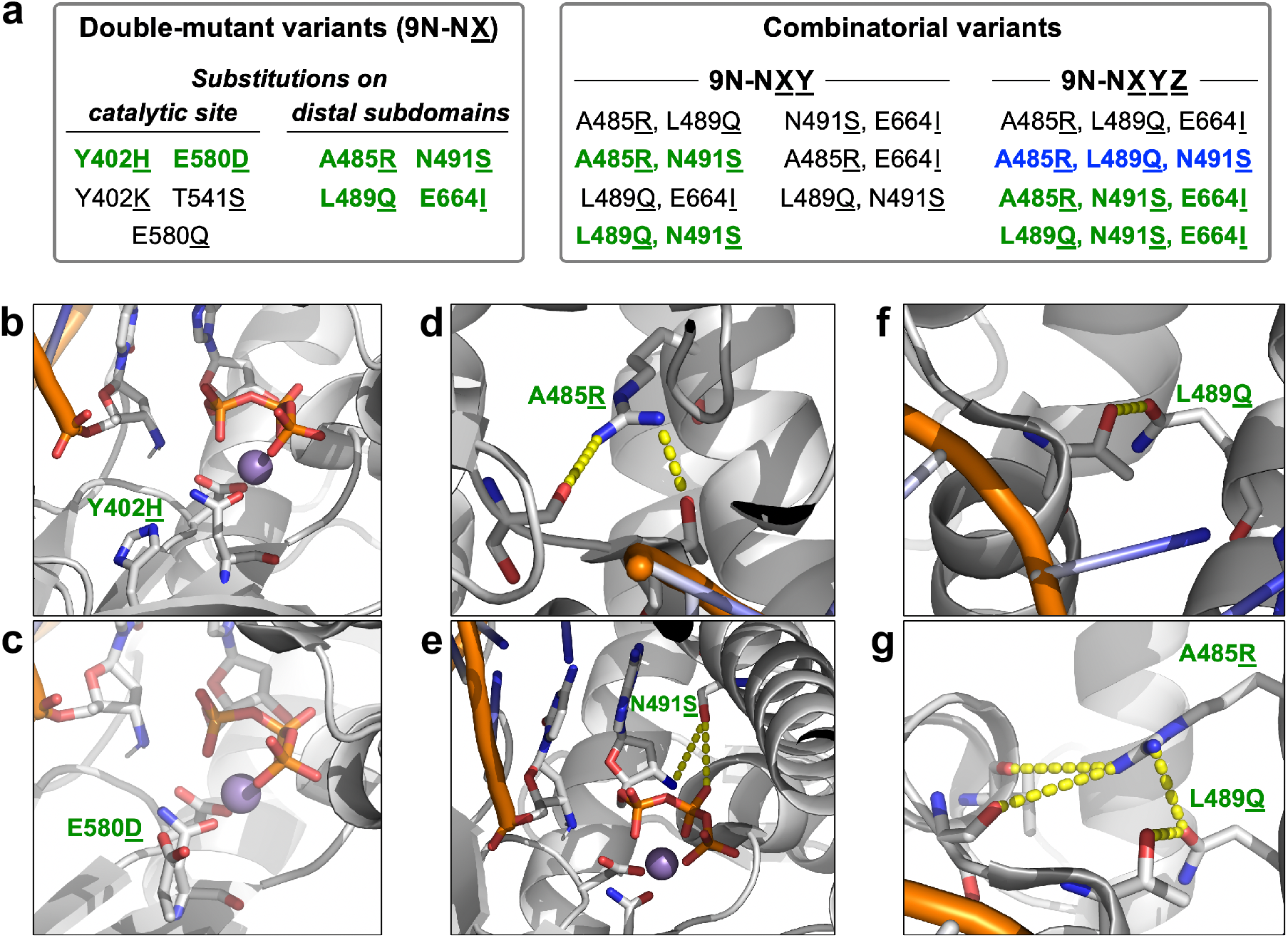
Rosetta modeling-guided double and combinatorial mutagenesis of 9°N-N. **(a)** Classification of variants based on site of amino acid substitution and combinatorial status. Substitutions leading to an increase in NP-DNA polymerase activity are listed in green, the variant with the highest catalytic rate is highlighted in blue. **(b–g)** Rosetta-based structural models of selected key substitutions, highlighting the second-shell effects on the catalytic site (**b** and **c**) and the distal finger subdomain (**d**–**g**).

First, we sought to reduce divalent metal occupancy in the active site by altering the hydrogen bond network: Y402H (9°N-NH) (**Figure 3b**) and E580D (9°N-ND) (**Figure 3c**). Further remodeling was conducted to increase local positive charge: Y402K (9°N-NK) and E580Q (9°N-NQ**). We also investigated making a stronger hydrogen bond with the primer 3’-amino group: T541S (9°N-NS*). The substitutions in 9°N-NH and 9°N-ND resulted in ~2.5-fold primer-extension rate enhancements (**Table S4**), however for the other three variants, the rates were slower than that for 9°N-N by ~20%.

Certain processive mutations in KOD, an ortholog of 9°N, have been reported to increase the polymerization rate of threose nucleic acid,^30^ which is another non-natural nucleic acid polymer that adopts a helical geometry similar to that of A-form RNA. We therefore sought to transfer some of these mutations to 9°N-N to facilitate a faster NP-DNA extension. To this end, we modeled four double-mutant variants with substitutions on the distal subdomains: A485R (9°N-NR) (**Figure 3d**), N491S (9°N-NS) (**Figure 3e**), L489Q (9°N-NQ) (**Figure 3f**), and E664I (9°N-NI). In our models, the finger subdomain substitutions A485R and L489Q formed additional hydrogen bonds with the residues S348 and T349, which are at the N-terminus of a helix (residues 349–359). This N-terminal segment in turn makes hydrogen bonding and electrostatic interactions (via its helix dipole) with the downstream 5’-phosphate of the DNA template. These interactions may thus indirectly help stabilize the template in an orientation productive for the catalysis. Furthermore, position 491 is located on the active-site-facing side of the helix, and in our model, S491 is poised to make hydrogen-bonding interactions with both the pyrophosphate and 3’-amino group of the incoming monomer. Based on these possible favorable interactions, we synthesized these double-mutant variants. Gratifyingly, these variants resulted in 2.6- to 3.3-fold primer-extension rate enhancements.

Active-site and distal substitutions that led to an increase in catalytic activity were then investigated for synergistic effects through the synthesis of triple-mutant variants (9°N-NRQ, 9°N-NRS, 9°N-NQI, 9°N-NQS, 9°N-NSI, 9°N-NRI, and 9°N-RQS) and quadruple-mutant variants (9°N-NQSI, 9°N-NRQS, 9°N-NRSI, and 9°N-NRQI). Among the triple-mutant variants, 9°N-NRS and 9°N-NQS provided significantly larger (~10-fold) *k*_obs_. Majority of the quadruple-mutant variants had a similar activity to 9°N-NRS and 9°N-NQS, while 9°N-NRQS (**Figure 3g**) catalyzed N3→P5’ phosphoramidate bond formation with the fastest rate (*k*_obs_ = 0.13 h^−1^). Of note, compared to the extension kinetics for double-mutant variants, *k*_obs_ for 9°N-NRS and 9°N-NQS were ~4-fold faster, whereas *k*_obs_ for 9°N-NRQI was 1.4-fold slower. These findings suggest that N491S substitution, in combination with A485R and L489Q substitutions, can synergistically improve the polymerase processivity.

### Assessing NP-DNA and DNA Polymerase Activity

To shed light on the factors governing polymerase activity, we examined the enzyme dependence on divalent metal ion and 3’-nucleophile (**Figure 4a**). For these studies, we focused on 9°N-N and 9°N-NRQS, as the former variant constitutes the amino acid substitution essential for the desired unnatural activity and the latter produces internucleotidyl phosphoramidate linkages at fastest rates among all 9°N variants we screened. For both 9°N-N and 9°N-NRQS, the primer extensions, whether the DNA primer has a 3’-OH or 3’-NH2 nucleophile, were substantially faster with Mn^2+^ than Mg^2+^. For 9°N-N, *k*_obs_ of the 3’-amino-primer extension with 3’-NH_2_-ddGTP in the presence of Mn^2+^ was 0.01 h^-1^ (**Figure 4b**, **entry 8**). 9°N-NRQS catalyzed the formation of phosphoramidate bonds with ~13-fold enhanced kinetics at 55 °C compared to 9°N-N (**entry 12**). Further investigations on polymerase activity revealed that substituting Asp-404 with Asn-404 (D404N) was necessary for catalyzing the formation of N3→P5’ phosphoramidate bonds: 9°N-RQS (**Table S1** and **Figure S2C**), which is the close analog of 9°N-NRQS with Asp-404 intact, lost this unnatural catalytic function completely.

**Figure 4.**
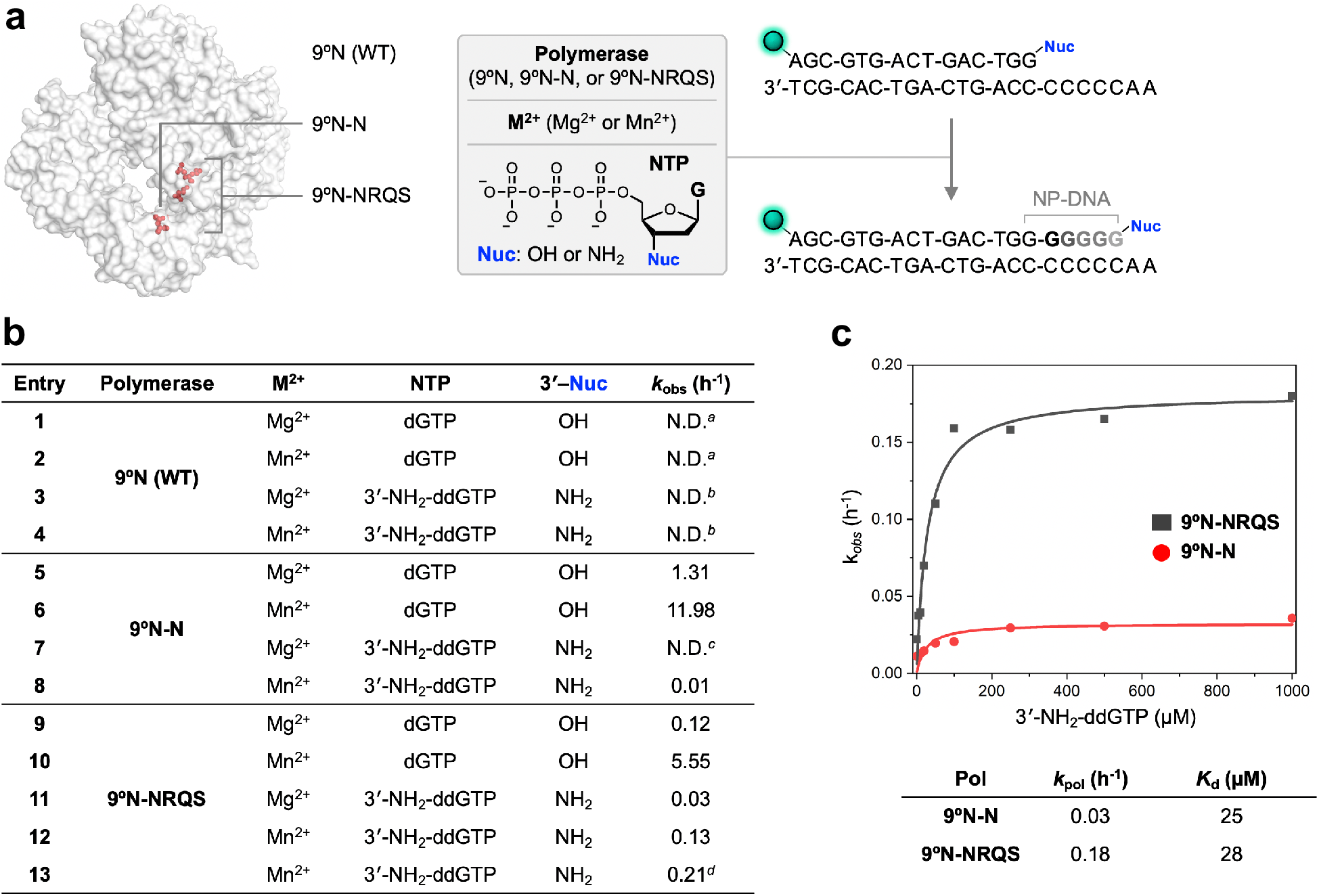
Enzymatic DNA and NP-DNA primer-extensions. **(a)** Primer-extension reaction conditions. 5’ end of the primer strand is labeled with 6-carboxyfluorescein (FAM), represented as green sphere. **(b)** Primer-extension reaction rates. *a*: Product formation is complete in less than 1 min; *b:* Extension product is undetectable; *c*: Extension product is detectable, but the rate of formation is too slow to measure; *d:* Reaction temperature is 65 °C. **(c)** Comparison of maximum rates of 3’-amino-G incorporation, *k*_pol_, and dissociation constants, *K_d_,* for both 9°N-N and 9°N-NRQS.

Given the thermophilic character of 9°N, we sought to explore how reaction temperature affects the NP-DNA polymerase activity of 9°N-NRQS. Reaction mixtures with identical contents (3’-amino-G primer/5-A_2_C_5_ DNA template, 3’-NH_2_-ddGTP, Mn^2+^) were incubated for 12 h at temperatures ranging from 45 to 75 °C with 5 °C intervals (**Figure S4**). The combined yields of the extension products were lowest (32–34% for the total of +1-nt and +2-nt products) for the samples incubated at temperatures below 55 °C. A similar outcome was observed for the 75 °C sample (52% for the total of +1-nt, +2-nt, and +3-nt products), likely due to the irreversible destabilization of the primer/template complex or inactivation of the polymerase at this temperature. Efficiency of the phosphoramidate bond formation was highest at 65 and 70 °C, with the +3-nt product fraction reaching up to ~17%. Due to possibility of the higher risk of polymerase inactivation for longer incubations at 70 °C, we elected 65 °C as the optimum reaction temperature. At 65 °C, *k*_obs_ of 3’-amino-G primer extension with 3’-NH2-ddGTP was 0.21 h^−1^ (**Figure 4b**, **entry 13**), a 60% enhancement of the kinetics observed for the 55 °C condition. This primer-extension rate was similar to that we measured for a longer primer/template complex. A 21-nt primer on full length complementary DNA with the same overhang (5’-A_2_C_5_) displayed *k_obs_* of 0.18 h^−1^, suggesting that our 15-nt base-pair primer/template duplex does not suffer from thermal destabilization at 65 °C.

We compared the NP-DNA polymerase activities of 9°N-N and 9°N-NRQS by determining the maximum rate of 3’-amino-G incorporation, *k*_pol_, and dissociation constant, *K*_d_ (**Figure 4c**). For 9°N-N and 9°N-NRQS, *k*_pol_ rate constants were 0.03 h^−1^ and 0.18 h^−1^, respectively, which was 6-fold increase in catalytic efficiency. *K*_d_ of 3’-NH_2_-ddGTP with 9°N-N and 9°N-NRQS were 25 μM and 28 μM, respectively, suggesting the similar binding affinity of 3’-NH_2_-ddGTP monomer with both enzymes.

As the 9°N active-site mutations enable enzymatic NP-DNA synthesis, we sought to gain insight into how these mutations affect DNA polymerase activity. Both 9°N-N and 9°N-NRQS displayed significantly reduced activity for extending the canonical 3’-hydroxyl DNA primer with dGTP. D404N substitution, not surprisingly, appeared to be the major cause of the reduced extension rate. When 9°N-N was used as the polymerase, *k*_obs_ values were measured as 1.3 h^−1^ for Mg^2+^ (**Figure 4b**, **entry 5**) and 11.98 h^−1^ for Mn^2+^ (**entry 6**). DNA extension was slowest (*k*_obs_ = 0.12 h^−1^) with 9°N-NRQS and Mg^2+^ at 55 °C (**entry 9**).

### Primer-Extension Kinetics with Homopolymeric DNA Templates

Next, we sought to compare the 3’-amino-primer extensions on homopolymeric DNA templates with other 3’-NH_2_-ddNTPs (**Figure 5a**). Monomers 3’-NH_2_-ddTTP, 3’-NH_2_-ddATP, and 3’-NH_2_-ddCTP were incubated separately with mixtures containing DNA templates 5’-C_2_T_5_, 5’-C_2_A_5_, and 5’-T2G5, respectively (**Figure 5b**), under the reaction conditions described above for 3’-NH_2_-ddGTP. The rate of primer extension on 5’-C_2_T_5_ with 3’-NH_2_-ddATP (*k*_obs_ = 0.20 h^−1^) was similar to that on 5’-A2C5 with 3’-NH_2_-ddGTP (*k*_obs_ = 0.21 h^−1^) (**Figure 5c**). The rates of primer-extension on 5’-C_2_A_5_ with 3’-NH_2_-ddTTP and on 5’-T2G5 with 3’-NH_2_-ddCTP were slightly slower (*k*_obs_ = 0.16 h^−1^ and 0.15 h^−1^, respectively). Despite nearly 90% or more of the primer being consumed within 24 hours, the +3-nt product band intensities for the reactions involving 3’-NH_2_-ddCTP and 3’-NH_2_-ddTTP plateaued at 23% and 5%, respectively. The extension reactions involving 3’-NH_2_-ddGTP and 3’-NH_2_-ddATP proved to be fastest: the +3nt products were observed within 3 hours, with the band intensities reaching to 37% and 12%, respectively, after 24 hours. Within this timeframe, the primer extension with 3’-NH_2_-ddGTP provided +4-nt product, albeit at 5% yield. This finding suggests that the primer-extension rate is substantially influenced by factors other than canonical nucleobase hydrogen bonding between the 3’-NH_2_-ddNTPs and the template within the 9°N-NRQS binding site. Such factors, including base stacking and sterics, have been recognized to play critical roles in governing enzymatic nucleic acid polymerization.^31^

**Figure 5.**
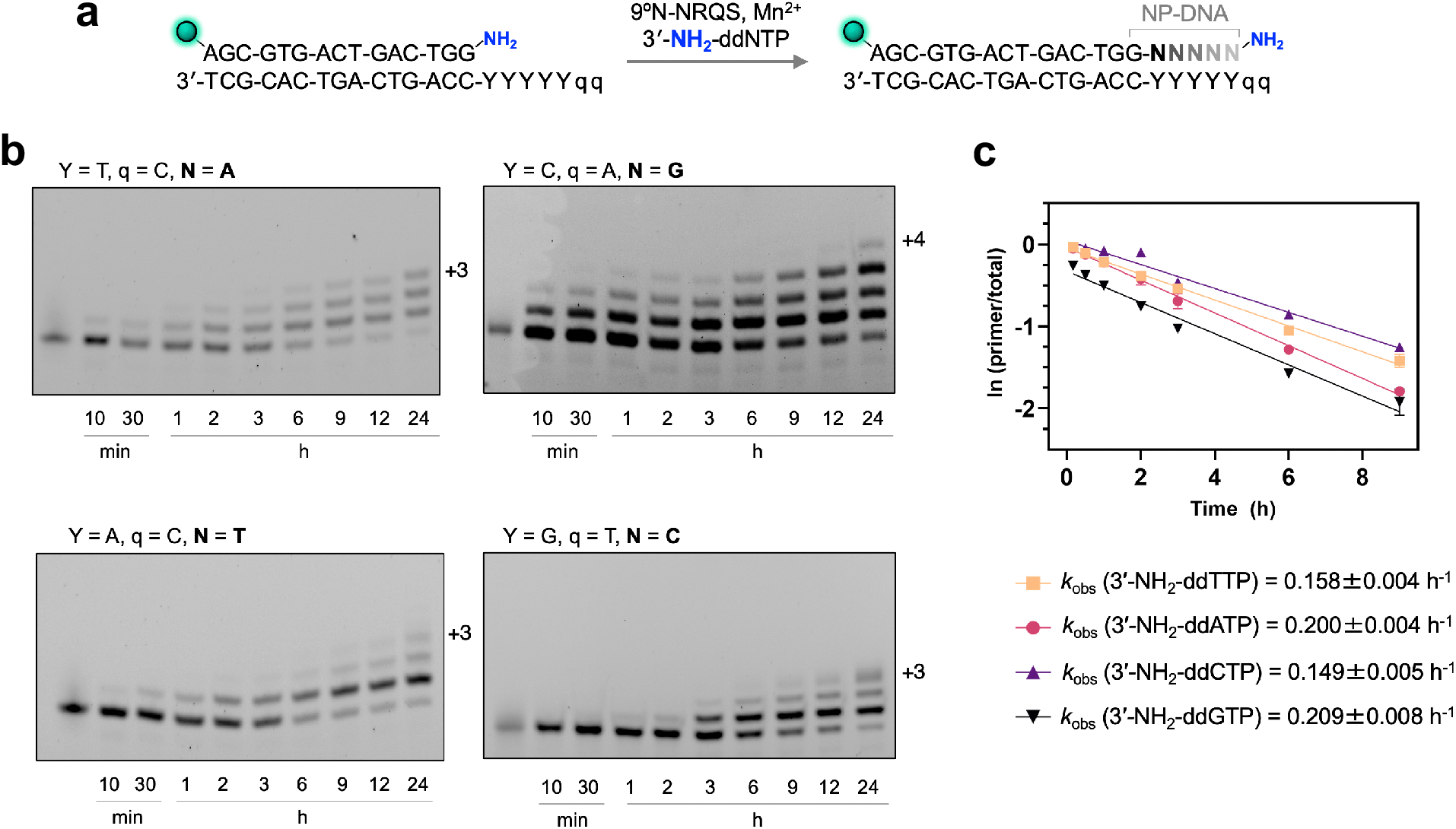
9°N-NRQS-catalyzed homopolymeric NP-DNA primer-extensions on a DNA template. **(a)** Primer-extension reaction condition. Four different reaction mixtures were prepared using a specific 3-NH_2_-ddNTP (N: base incorporated to the primer). Y: template base; q: overhang. **(b)** Denaturing SDS-PAGE analyses of the extension reaction mixtures. Imaging performed at FAM excitation (epi-blue, 460–490 nm). **(c)** (Top) natural log of the fraction of the unreacted primer strand versus incubation time, (bottom) observed rate constants, *k*_obs_. Error estimates were calculated from the standard deviation of three replicates.

### Template-Directed Polymerase Activity

The DNA template-dependent activity of 9°N-NRQS was assessed using polymerase end point assays. By comparing the 3’-amino-primer extensions from a total of sixteen nucleotide-template combinations at both 55 and 65 °C, we determined the rate of nucleotide-template mismatch with respect to the Watson-Crick base pairing (**Figure 6a**). The buffered mixtures of 9°N-NRQS, Mn^2+^, the 3’-amino-G primer, and a DNA template selected from 5’-C_2_A_5_, 5’-C_2_T_5_, 5’-T_2_G_5_, and 5’-A_2_C_5_ were incubated with all four 3’-NH_2_-ddNTPs. Mixtures were quenched after 24 h and percentages of the primer and the extension product fractions were measured after denaturing PAGE separation. For each nucleotide-template combination, the nucleotide incorporation fidelity was expressed as discrimination ratio (*D*), which is defined as [Watson-Crick extension product] / [total mismatch product]. For most of the samples incubated at 55 °C, *D*s were above 10, displaying at least an order of magnitude nucleotide selectivity. The degree of mismatch was highest for 3’-NH_2_-ddTTP: The *D* values were determined as 2.9, 1.1, and 2.9 for 5’-C_2_T_5_, 5’-T_2_G_5_, and 5’-A_2_C_5_ templates, respectively. It should be noted that these *D*s represent the upper bound of mutation rate, as the mismatch conditions included a single type of 3’-NH_2_-ddNTP. The 3’-amino-G primer was prone to a higher degree of nucleotide misincorporation when the temperature was increased. For the reaction mixtures incubated at 65 °C, *D*s decreased by 5- to 10-fold compared to those incubated at 55 °C.

**Figure 6.**
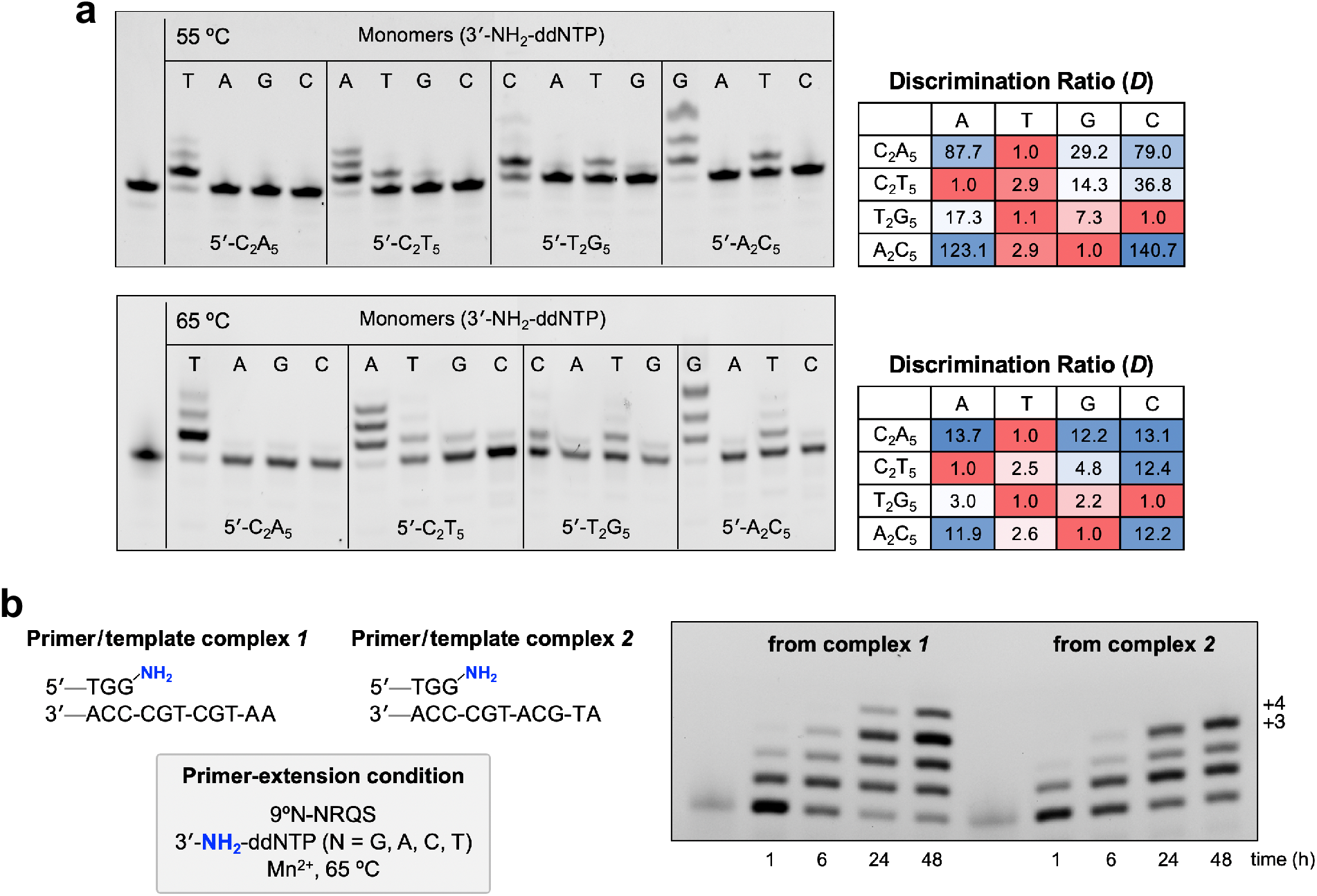
Effect of Template on 9°N-NRQS-catalyzed NP-DNA primer-extension. **(a)** Investigation of the NP-DNA polymerase activity using the nucleotide-template mismatch assays at 55 °C (top) and 65 °C (bottom). **(b)** Investigation of the NP-DNA polymerase activity of 9°N-NRQS on templates with mixed sequences.

Finally, we investigated NP-DNA polymerase activity of 9°N-NRQS with templates containing mixed sequences (**Figure 6b**). The extension of the same 15-nt 3’-amino-G primer on DNA templates 5’-AATGCTGC and 5’-ATGCATGC in the presence of all 3’-NH_2_-ddNTPs were examined. The comparison of these two templates was intended to gain insight into the effect of addition of T on the primer-extension efficiency. Within 24 hours of incubation at 65 °C, the +4-nt and +3-nt products were observed for the mixture including templates 5’-AATGCTGC and 5’-ATGCATGC, respectively. At the 48-hour mark, the relative abundance of these extension products reached up to 15% and 34%. Although the addition rates of all 3’-NH_2_-ddNTPs are modest, replacing T with A on the template resulted in a lower extension efficiency, thus a shorter extension product. Furthermore, the absence of longer products (≥ +5-nt) suggests that either 9°N-NRQS loses catalytic activity drastically after 24 hours of incubation or the helical geometry of the extending primer-template duplex displays stalling effect on the enzyme. Although either proposition may be correct, no major extension beyond four consecutive N3’→P5’ phosphoramidate linkages was observed using a 16-nt primer containing a single phosphoramidate bond (**Table S1**).

## DISCUSSION

WT polymerases have evolved to catalyze the formation of internucleotidyl phosphodiester linkages in genetic polymers. 9°N, an archaeal B-family DNA polymerase, can accommodate non-canonical deoxynucleoside triphosphate monomers for extending primers but does not catalyze the formation of internucleotidyl phosphoramidate linkages. We hypothesized that the 3’-amino group of a primer could serve as a nucleophile in an engineered 9°N active site with altered metal-binding properties while maintaining the sterics required to accommodate the nucleophile. Through a mechanism-guided structure-based design approach, we performed systematic active-site mutagenesis to transform 9°N into a catalyst that can slowly extend 3’-amino terminated primers with 3’-NH_2_-ddNTP monomers.

Notably, the formation of the N3’→P5’ phosphoramidate bond was observed when Asp-404, one of the two metal-binding Asp residues in the active site, was replaced with Asn (9°N-N). This suggests that an active-site residue with little affinity towards a divalent metal but more polar than aliphatic or aromatic counterparts could allow phosphorylation of a primer 3’-amino group. Further metal-binding-site remodeling aimed at altering the electrostatics, hydrogen bonding, and hydrophobicity of the catalytic site by additional single substitutions resulted in additional, albeit marginal, increases in activity. These results highlight the possibilities as well as limits on repurposing a highly evolved active site for a promiscuous activity by solely focusing on screening mutations in the catalytic site. As enzyme activity and selectivity likely emerge by the collective action of many protein structural elements, subsequent exploration of substitutions was focused on the distal subdomains of the polymerase, which were expected to be key elements underlying processivity of 9°N based on previous studies with KOD.^30^ Substitutions A485R, L489Q, and N491S provided 9°N-NRQS as the most active variant (among all 26 variants expressed), with up to ~13-fold enhanced kinetics at 55 °C compared to 9°N-N. Investigation of the effect of temperature on primer-extension kinetics revealed that 9°N-NRQS can catalyze the formation of the N3’→P5’ phosphoramidate bond, at least during the first 12 hours of incubation, at elevated temperatures. At 65 °C, 9°N-NRQS produced phosphoramidate bonds with a rate further enhanced by nearly 60%, which corresponds to a ~21-fold enhanced rate compared to that measured with 9°N-N at 55 °C. The *k*obs values were measured as 0.2 h^−1^ for the extensions of the 3’-amino-G primer on 5’-A_2_C_5_ and 5’-C_2_T_5_ DNA templates with 3’-NH_2_-ddGTP and 3’-NH_2_-ddATP, respectively. Taken together, these results show that 9°N-NRQS can extend 3’-amino-G primer on DNA templates at rates similar those measured for non-enzymatic template-directed extensions with the highly activated 3’-amino-nucleoside phosphoroimidazolides^19^ and indicate how catalytic-site distal elements may also play a key role in the proper positioning of substrates leading to a significant enhancement of the promiscuous activity of the enzyme. Further improvement in the 3’-amino primer extension rate is necessary to make 9°N, KOD, or other ortholog polymerases viable biocatalysts for the synthesis of long NP-DNA sequences, which may be achieved through directed evolution.

It is crucial to note that 9°N-NRQS has substantially reduced polymerase activity for extending the canonical 3’-hydroxyl DNA primer. DNA extension from dGTP using 9°N-NRQS is slowest in the presence of Mg^2+^ at 55 °C. As a consequence, this variant displays an unnatural polymerase activity (catalyzing NP-DNA extension) similar to what was a natural activity (catalyzing DNA extension) prior to the active-site mutations. It is interesting to note that unlike examples of evolutionary trajectories in natural enzymes,^32^ there is a clear trade-off between the original and designed promiscuous activities of 9°N. Such a precipitous trade-off was previously observed between deaminase (original) and organophosphate hydrolase (designed) activities in another redesigned metalloenzyme.^22^ These results suggest that the highly tuned environment in metal binding sites may also be highly susceptible to substitution of metal-chelating residues, and can provide a general strategy to explore novel activities in metalloenzymes by redesign.

Non-natural nucleic acid polymers are emerging tools with major potential in synthetic biology, biotechnology, and molecular medicine.^33–36^ Recent reports showed that these polymers, whether in homopolymeric or hybrid form, can store and transfer genetic information^11,37–39^ as well as evolve to bind a molecular target^40–43^ or catalyze a reaction.^44^ In addition to gaining insights on the catalytic mechanisms and latent promiscuity in enzymes, we also expect that new polymerase functions, combined with the expansion of genetic information space, will continue to push the boundaries of designer biological systems.

## ACKNOWLEDGMENTS

We thank Dr. John C. Chaput for his generous donation of the 9°N-WT plasmid. This work was supported by the American Cancer Society, Institutional Research Grant Early Investigator Award and the Rutgers Cancer Institute of New Jersey NCI Cancer Center Support Grant (P30CA072720) and US National Institutes of Health / NIBIB Trailblazer Award (R21 EB029548) (to E.C.I.), National Science Foundation (NSF) grant CBET-1929237 and National Institutes of Health (NIH) grant GM132565 (to S.D.K.).

## AUTHOR CONTRIBUTIONS

S.D.K. and E.C.I. conceived the study. T.A. and W.A.H. conducted structure-guided enzyme redesign. T.A. performed protein expression and isolation. T.A. and J.H. performed primer-extension experiments. A.G. conducted MD simulations. All of the authors contributed to the interpretation of data. T.A., S.D.K., and E.C.I. wrote the manuscript with input from all authors. D.M.Y., S.D.K., and E.C.I. supervised the study.

