## Supplementary Information for "Introducing a New Bond-Forming Activity in an Archaeal DNA Polymerase by Structure-Guided Enzyme Redesign"

#### Contents




| | |
|---|---|
| **Chemicals, Materials, and Biologics** | S3–S4 |
| **Divalent Metal Screening for 9ºN-WT and 9ºN-N** | S5 |
| **Computational Modeling Using Rosetta** | S6 |
| **Protein Expression and Purification** | S7–S10 |
| **Primer Extension Assays and Kinetics** | S11 |
| **Comparative Polymerase Activity Analysis** | S12 |
| **Effect of Temperature on the NP-DNA Pol Activity of 9ºN-NRQS** | S13 |
| **References** | S14 |

#### Chemicals, Materials, and Biologics

**Reagents:** DNA oligonucleotides (**Table S1**) were purchased from Sigma Life Sciences. Therminator DNA polymerase and 2′-deoxyribonucleotide 5′-triphosphates were purchased from New England Biolabs. 3′-Amino-2′,3′-dideoxyribonucleotide 5′-triphosphates were obtained from TriLink BioTechnologies. Ampicillin (Amp), Luria broth (LB), isopropyl-β-D-thiogalactopyranoside (IPTG), sodium chloride (NaCl), ammonium sulfate ($(NH_4)_2SO_4$), hydrochloric acid (HCl), glycerol, imidazole, glycerol, tris(hydroxymethyl)-aminomethane (Tris), potassium chloride (KCl), ethylenediaminetetraacetic acid (EDTA), and urea were purchased from Millipore-Sigma. sodium hydroxide (NaOH) was purchased from Fisher Scientific Inc. Polyethylenimine (PEI) was purchased from BeanTown Chemical.

Buffers were prepared freshly in Milli-Q® water and their pH was adjusted using HCl or NaOH (pH 8.8 for ThermoPol buffer and pH 8.0 for all the other buffers). All solutions were autoclaved or sterile filtered (0.2 µm) prior to use.

Lysis buffer: Tris (10 mM), NaCl (500 mM), EDTA (100 µM), glycerol (10% v/v).

Wash buffer: Tris (10 mM), NaCl (600 mM), imidazole (20 mM), glycerol (10% v/v).

Elution buffer: Tris (10 mM), NaCl (600 mM), imidazole (500 mM), glycerol (10% v/v).

Dialysis buffer: Tris (10 mM), NaCl (600 mM), EDTA (500 µM), glycerol (10% v/v).

Polymerase storage buffer: Tris (10 mM), NaCl (200 mM), glycerol (10% v/v).

ThermoPol buffer (1x): Tris (20 mM), $(NH_4)_2SO_4$ (10 mM), KCl (10 mM), 0.1% Triton X-100.

Stop buffer: EDTA (10 mM) and urea (6 M).

TE buffer: Tris (10 mM), EDTA (1 mM).

**Materials:** Ni-NTA sepharose affinity resin was purchased from Qiagen. Petri dish, culture tube, Amicon filter, dialysis tube and Falcon tubes were obtained from VWR international. PD-10 columns were purchased from GE Scientific.

**Biologics:** BL21-DE3 and XL10-Gold competent *E. Coli* cells were obtained from New England Biolabs. Plasmid of 9ºN-WT was obtained from the Chaput Lab (University of California, Irvine).

**Table S1.** List of oligonucleotide sequences used for primer extension assays.

| Sequence Name | Sequence (5′→3′) |
|---|---|
| 3′-Hydroxyl (canonical) primer | FAM-AGCGTGACTGACTGG |
| 15-nt 3′-amino primer † | FAM-AGCGTGACTGACTGG-$NH_2$ |
| 16-nt 3′-amino primer ‡ | FAM-AGCGTGACTGACTGGG-$NH_2$ |
| 14-nt primer precursor | FAM-AGCGTGACTGACTG |
| $C_2A_5$ template | CCAAAAACCAGTCAGTCACGCT |
| $C_2T_5$ template | CCTTTTTCCAGTCAGTCACGCT |
| $A_2C_5$ template | AACCCCCCCAGTCAGTCACGCT |
| $T_2G_5$ template | TTGGGGGCCAGTCAGTCACGCT |
| 20-nt DNA reference | FAM-AGCGTGACTGACTGGTTTTT |

† 15-nt 3′-amino primer was synthesized from 14-nt precursor via addition of a single 3′-$NH_2$-ddGTP using 9ºN-WT. ‡ 16-nt 3′-amino primer was synthesized from 14-nt precursor via addition of two 3′-$NH_2$-ddGTPs using 9ºN-WT.

The 14-nt primer precursor (5 μM), 5′-$T_2G_5$ template (5 μM), 3′-$NH_2$-ddGTP (100 μM), $Mg^{+2}$ (1 mM) and 9ºN-WT (5 μM) was incubated in 1x ThermoPol buffer at 12 °C for 10 min. Reaction was quenched by addition of the stop buffer. Reaction contents were loaded on a 1.5 mm thick 20% polyacrylamide gel and presence of the +1-nt product (3′-amino primer) was verified by gel imaging. The 3′-amino primer band was excised using a clean scalpel, isolated gel slab was finely crushed, and the resulting particulates were transferred to a 15 mL tube. The 3′-amino primer was then eluted from the gel particulates using a previously reported freeze-thaw method.[1] After the elution process, the isolated filtrate was lyophilized overnight. Dried pellet was resuspended in 1 mL nuclease-free water, to which 100 μL sodium acetate (3 M, pH 5.5) followed by 2.2 mL ethanol were added. The resulting mixture was chilled at -20 °C overnight, and then centrifuged (20,000 rcf, 4 °C, 1 h) to precipitate the 3′-amino primer. The supernatant was removed, and the resulting pellet was washed with 70 % ethanol and centrifuged (20,000 rcf, 4 °C, 15 min). The supernatant was discarded, and the pellet was resuspended in nuclease-free water. The oligonucleotide concentration was determined by UV (NanoDrop™) measurements. Product purity was determined as 95% based on 20% denaturing PAGE analysis.

#### Divalent Metal Screening for 9ºN-WT and 9ºN-N

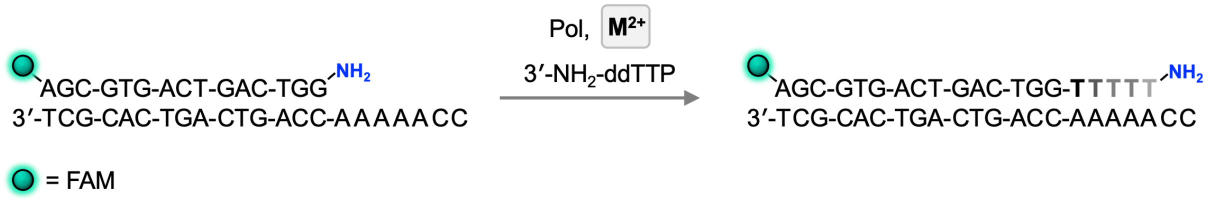


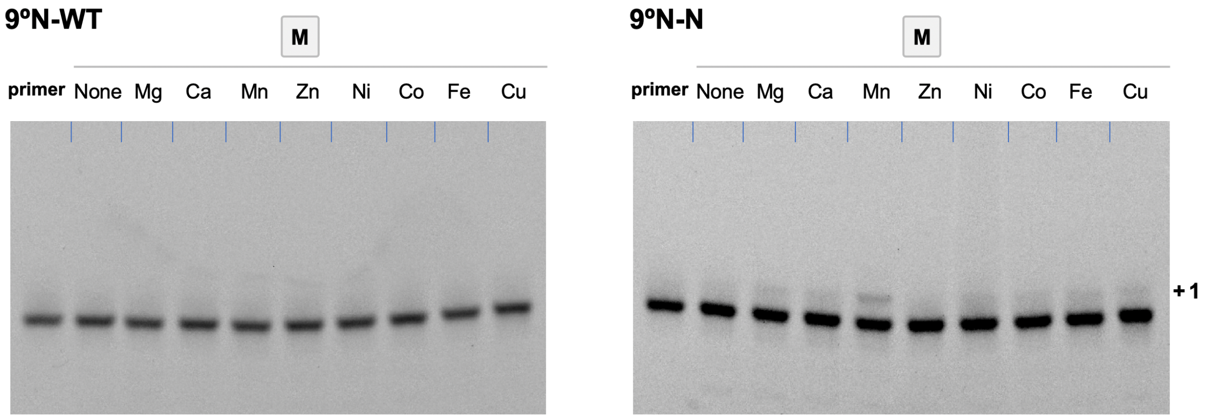


**Figure S1.** Divalent metal ion ($M^{2+}$) screen using 9ºN-WT and 9ºN-N. Reaction temperature: 55 ºC. Reaction time: 12 h.

#### Computational Modeling Using Rosetta

PyRosetta[2] was used to analyze the enzyme structure and perform in silico mutagenesis. To explore substitutions that were expected to alter the microenvironment of Asp404, we substituted all neighboring residues that are non-catalytic to all 19 amino acids. All residue pairs with Cα-Cα distance <5.5Å were considered neighbors, and residue pairs with Cα-Cα distance <11Å were also considered neighbors if their Cα-Cβ vectors were at an angle <75°.[3] Structural models for each variant were computed by replacing the reference side chain atomic coordinates in the starting model generated from the crystal structure of 9N polymerase (PDB code: 5OMV) with those of the substituted amino acid(s) and performing three rounds of Monte Carlo optimization of rotamers for all side chains (except those involved in catalysis) falling within an 8Å radius of the substitution(s), followed by gradient-based energy minimization of the entire structure, with atom positional restraints on the backbone, incoming monomer and the primer-template complex to limit significant changes to backbone geometry.[4] Computed structural model optimizations were performed with the REF2015.[5] Energetic consequences of amino acid substitutions were determined by performing identical side chain optimization and energy minimization on both wild-type and substituted models thrice and subtracting the total energy of the lowest-scoring wild-type model from that of the lowest-scoring substituted model.[6] Energy changes within 3 Rosetta Energy Units for each residue position were considered acceptable and the resulting models were visually inspected to choose substitutions for experimental characterization.

#### Protein Expression and Purification

Site-directed mutagenesis (SDM) and transformation: All the mutations in the plasmid were incorporated by SDM (see **Tables S2** and **S3**) and were sequence confirmed by commercial Sanger sequencing (Genwiz). Plasmids containing N-terminal His tag and Amp resistance gene were transformed into BL21(DE3) competent cells using heat shock method and were selected on the LB plate with Amp (100 µg/mL).

Starter culture and IPTG induction: A 5 mL of LB containing Amp (100 µg/mL) was inoculated with a single isolated colony and was grown at 37 °C overnight. Next day, the overnight culture was used to inoculate the 500 mL of LB with Amp (100 µg/mL) and grown at 37 °C until the optical density ($OD_{600}$) of culture reached 0.4-0.6. Protein expression was induced by the addition of IPTG to a final concentration of 500 µM. The sample was shaken at 18 °C for a minimum of 18 h.

Protein purification: Purification of the 9°N variants was performed based on a previously reported protocol[7] with modifications. The cells were harvested by centrifugation (4000 x g, 30 min, 4 °C) and resuspended in 30 mL of lysis buffer. Cells were lysed by sonication for 10 min on ice. Cell lysate was incubated first in a 80 °C water bath for 60 min to denature the endogenous *E. Coli* proteins and then on ice for 30 min. Cell debris and endogenous proteins were removed by centrifugation (50,000 x g, 4 °C, 30 min). To remove the nucleic acids, 10% PEI was added to a final concentration of 0.5% PEI. The solution was incubated on ice for 15 min and centrifuged (50,000 x g, 4 °C, 30 min). To remove the excess PEI and precipitate the protein, ammonium sulphate was added to a final concentration of 60%. Protein was precipitated (50,000 x g, 4 °C, 30 min) and pellet was resuspended in wash buffer and filtered through a 0.22 µm syringe filter. Supernatant was then added to Ni-NTA gravity column which was pre-equilibrated with the wash buffer and manually eluted with the elution buffer (~15 mL). The eluted protein was dialyzed overnight into the buffer containing 0.5 mM EDTA to remove divalent metal ions to get apo-protein. The dialyzed protein buffer was exchanged with the polymerase storage buffer using a PD10 column and the isolated fraction was concentrated to 10 µM using a 30 kDa cutoff Amicon centrifugal filter (Millipore). The protein expression was validated by SDS-PAGE gel analysis (see **Figure S2**).

**Table S2.** Classification of 9ºN variants.

| Type of Variant | Name of Variant | Site of Substitution |
|---|---|---|
| Single | 9ºN-N | D404N |
| | 9ºN-A′ | D542A |
| | 9ºN-S | D542S |
| | 9ºN-T | D542T |
| | 9ºN-H | D542H |
| Double | 9ºN-NN′ | D404N, D542N |
| | 9ºN-NR | D404N, A485R |
| | 9ºN-NQ | D404N, L489Q |
| | 9ºN-NS | D404N, N491S |
| | 9ºN-NI | D404N, E664I |
| | 9ºN-NH | D404N, Y402H |
| | 9ºN-NK | D404N, Y402K |
| | 9ºN-NS* | D404N, T541S |
| | 9ºN-ND | D404N, E580D |
| | 9ºN-NQ** | D404N, E580Q |
| Triple | 9ºN-NRQ | D404N, A485R, L489Q |
| | 9ºN-NRS | D404N, A485R, N491S |
| | 9ºN-NQI | D404N, L489Q, E664I |
| | 9ºN-NQS | D404N, L489Q, N491S |
| | 9ºN-NSI | D404N, N491S, E664I |
| | 9ºN-NRI | D404N, A485R, E664I |
| | 9ºN-RQS | A485R, L489Q, N491S |
| Quadruple | 9ºN-NQSI | D404N, L489Q, N491S, E664I |
| | 9ºN-NRQS | D404N, A485R, L489Q, N491S |
| | 9ºN-NRSI | D404N, A485R, N491S, E664I |
| | 9ºN-NRQI | D404N, A485R, L489Q, E664I |

(′), (*), and (**) denote substitution of Asp-542, Thr-541, and Glu-580, respectively.

**Table S3.** List of oligonucleotide sequences used for protein expression. Listed: Forward (fwd) and reverse (rvs) primers for SDM experiments.

| Sequence Name | Sequence (5′→3′) |
|---|---|
| D404N fwd | GGATAACATTGTGTATCTGAATTTTCGTAGCCTGTATCCG |
| D404N rvs | CGGATACAGGCTACGAAAATTCAGATACACAATGTTATCC |
| D542N fwd | GTATGCGGATACCAATGGCCTGCATGC |
| D542N rvs | GCATGCAGGCCATTGGTATCCGCATAC |
| D404A fwd | GATAACATTGTGTATCTGGCTTTTCGTAGCCTGTATCC |
| D404A rvs | GGATACAGGCTACGAAAAGCCAGATACACAATGTTATC |
| D542A fwd | GTGCTGTATGCGGATACCGCTGGCCTGCATGCGACCATTC |
| D542A rvs | GAATGGTCGCATGCAGGCCAGCGGTATCCGCATACAGCAC |
| D542T fwd | CTGTATGCGGATACCACTGGCCTGCATGCGAC |
| D542T rvs | GTCGCATGCAGGCCAGTGGTATCCGCATACAG |
| D542S fwd | CTGTATGCGGATACCAGTGGCCTGCATGCGAC |
| D542S rvs | GTCGCATGCAGGCCACTGGTATCCGCATACAG |
| D542H fwd | GTATGCGGATACCCATGGCCTGCATG |
| D542H rvs | CATGCAGGCCATGGGTATCCGCATAC |
| A485R fwd | CTGGATTATCGTCAGCGCCGTATTAAAATTCTGGCCAAC |
| A485R rvs | GTTGGCCAGAATTTTAATACGGCGCTGACGATAATCCAG |
| E664I fwd | CGGAAAAACTGGTGATTCATATTCAAATTACCCGTGATCTGCG |
| E664I rvs | CGCAGATCACGGGTAATTTGAATATGAATCACCAGTTTTTCCG |
| N491S fwd | GCGATTAAAATTCTGGCCAGCAGCTTCTATGGCTATTATG |
| N491S rvs | CATAATAGCCATAGAAGCTGCTGGCCAGAATTTTAATCGC |
| L489Q fwd | CGCGCGATTAAAATTCAGGCCAACAGCTTCTATG |
| L489Q rvs | CATAGAAGCTGTTGGCCTGAATTTTAATCGCGCG |
| D404NA485RL489Q fwd | CGCCGTATTAAAATTCAGGCCAACAGCTTCTATG |
| D404NA485RL489Q rvs | CATAGAAGCTGTTGGCCTGAATTTTAATACGGCG |
| D404NA485RN491S fwd | CGTATTAAAATTCTGGCCAGCAGCTTCTATGGCTATTATG |
| D404NA485RN491S rvs | CATAATAGCCATAGAAGCTGCTGGCCAGAATTTTAATACG |
| D404NL489QN491S fwd | GCGATTAAAATTCAGGCCAGCAGCTTCTATGGCTATTATG |
| D404NL489QN491S rvs | CATAATAGCCATAGAAGCTGCTGGCCTGAATTTTAATCGC |
| D404NA485RL489QN491S fwd | CGTATTAAAATTCAGGCCAGCAGCTTCTATGGCTATTATG |
| D404NA485RL489QN491S rvs | CATAATAGCCATAGAAGCTGCTGGCCTGAATTTTAATACG |
| DALN491Y fwd | GCCGTATTAAAATTCAGGCCTACAGCTTCTATGGCTATTATG |
| DALN491Y rvs | CATAATAGCCATAGAAGCTGTAGGCCTGAATTTTAATACGGC |
| N404D fwd | GGATAACATTGTGTATCTGGATTTTCGTAGCCTGTATCC |
| N404D rvs | GGATACAGGCTACGAAAATCCAGATACACAATGTTATCC |
| 9ºN Sequencing primer | TGCGGATACCGATGGCCTGCATGC |

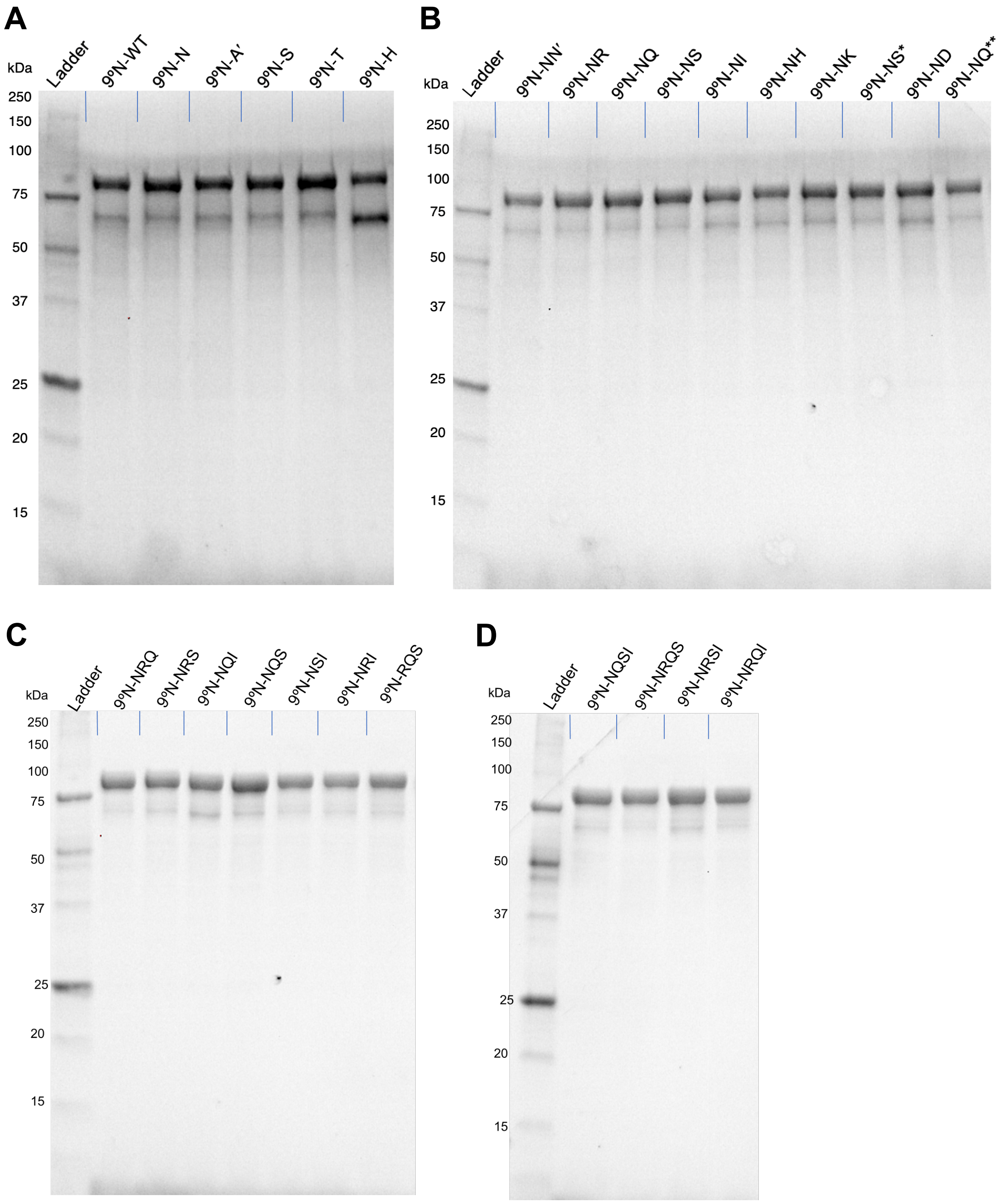


**Figure S2.** Analysis of polymerase isolation and purity by SDS-PAGE. **(A)** 9ºN single-mutant variants, **(B)** 9ºN double-mutant variants, **(C)** 9ºN triple-mutant variants, and **(D)** 9ºN quadruple-mutant variants. (′), (*), and (**) denote substitution of Asp-542, Thr-541, and Glu-580, respectively.

#### Primer Extension Assays and Kinetics

Prior to each primer-extension assay, the primer and template strands (1 μM each) and nucleoside triphosphate (100 μM) were dissolved in the ThermoPol buffer. The primer/template complex was formed by heating the mixture at 90 °C for 2 min and subsequently cooling to 4 °C with a ramp rate of 0.1 °C/s. The polymerase (1 μM) was then added to the cooled mixture. The primer-extension reaction was initiated by the addition of the divalent metal salt (1 mM). Reaction aliquots were taken at specific time intervals and quenched by adding the stop buffer. The resulting mixtures were then treated with formamide (50% v/v), heated at 80 °C for 2 min, and were subsequently separated by 20% denaturing PAGE with 7 M urea. Gels were analyzed on a BioRad Chemidoc MP gel imager and bands were quantified using Image lab software. The plots for enzyme kinetics were prepared using OriginLab software.

**Measurements of $k_{obs}$, $k_{pol}$, and $K_d$.** The first order reaction rate constant ($k_{obs}$) for primer-extension reaction was calculated by plotting the natural log of remaining primer versus time and was fitted to a linear equation.

For $k_{pol}$, and $K_d$ measurement[8], primer-extension assays were performed at various concentrations of 3′-$NH_2$-ddGTP (1, 10, 20, 50, 100, 250, 500, and 1000 μM) with 3′-amino-G primer (1 μM), 5′-$A_2C_5$ DNA template (1 μM), $Mn^{2+}$ (1 mM), and 9°N-NRQS (1 μM) in 1x ThermoPol buffer at 65 °C. Calculated $k_{obs}$ values for each of these reactions were then plotted against the concentration of 3′-$NH_2$-ddGTP and fitted according to the following equation:

$$k_{obs} = (k_{pol}\ [3'\text{-}NH_2\text{-ddGTP}] \,/\, K_d + [3'\text{-}NH_2\text{-ddGTP}])$$

, where $k_{pol}$ represents maximum rate of the nucleotide incorporation and $K_d$ represents dissociation constant.

#### Comparative Polymerase Activity Analysis

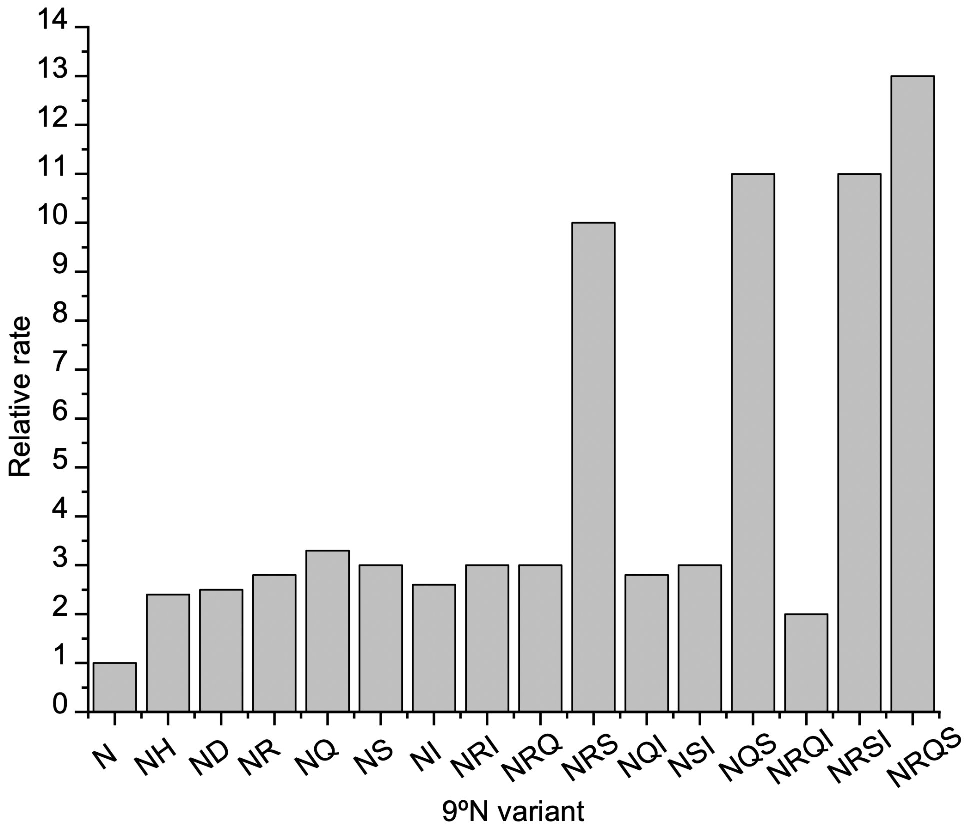


**Figure S3.** Relative rates of 3′-amino-G primer extension using productive 9ºN variants. The rates are normalized to that measured for 9ºN-N (labeled as "N"). Reactions were performed at 55 ºC and stopped after 12 hours. Reaction contents: 3′-Amino primer (1 μM), 5′-$A_2C_5$ template (1 μM), 3′-$NH_2$-ddGTP (100 μM), 9ºN variant (1 μM), $Mn^{2+}$ (1 mM), ThermoPol buffer (1x).

**Table S4.** Compilation of $k_{obs}$ for productive 9ºN variants.

| Variant | $k_{obs}$ ($h^{-1}$) | Relative rate | | Variant | $k_{obs}$ ($h^{-1}$) | Relative rate |
|---|---|---|---|---|---|---|
| N | 0.01 | 1.0 | | NRS | 0.10 | 10.0 |
| NH | 0.024 | 2.4 | | NQI | 0.028 | 2.8 |
| ND | 0.025 | 2.5 | | NSI | 0.03 | 3.0 |
| NR | 0.028 | 2.8 | | NQS | 0.11 | 11.0 |
| NQ | 0.033 | 3.3 | | NRQI | 0.02 | 2.0 |
| NS | 0.03 | 3.0 | | NRSI | 0.11 | 11.0 |
| NI | 0.026 | 2.6 | | NRQS | 0.13 | 13.0 |
| NRI | 0.03 | 3.0 | | NQSI | 0.10 | 10.0 |
| NRQ | 0.03 | 3.0 | | | | |

### Effect of Temperature on NP-DNA Pol Activity of 9ºN-NRQS

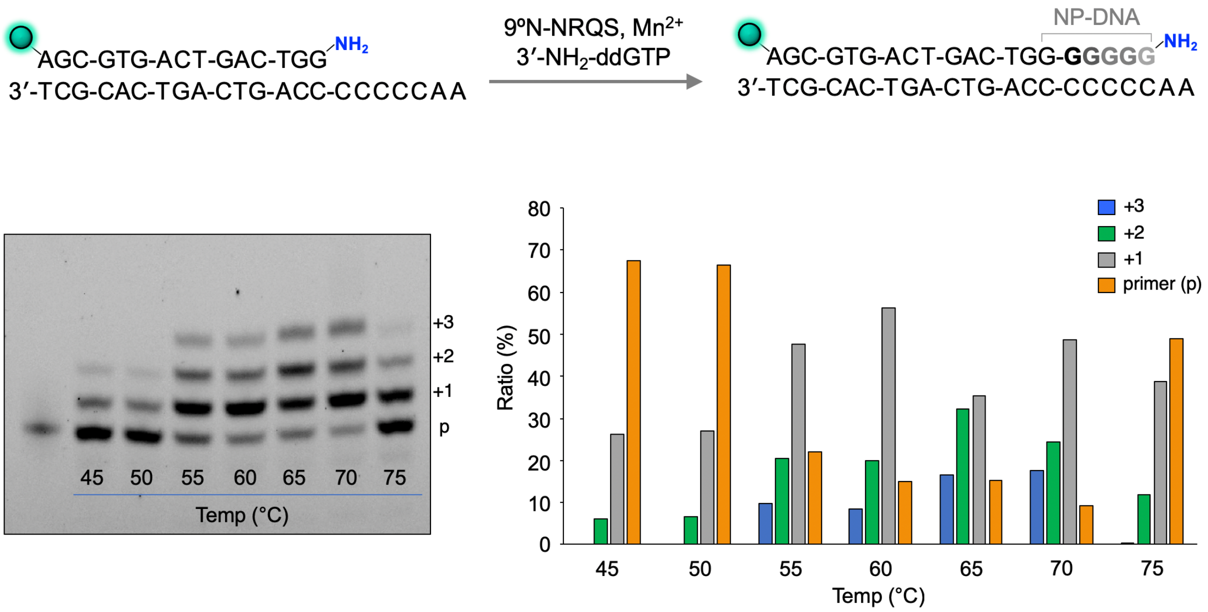


**Figure S4.** Effect of temperature on 9ºN-NRQS-catalyzed 3′-amino primer extension. Reaction time: 12 h.
